# Dynamics of coherent activity between cortical areas defines a two-stage process of selective attention

**DOI:** 10.1101/2020.02.09.940791

**Authors:** E. Levichkina, M. Kermani, Y.B. Saalmann, T.R. Vidyasagar

## Abstract

Analysing a visual scene requires the brain to briefly keep in memory potentially relevant parts and then direct attention to their locations for detailed processing. To reveal the neuronal basis of the underlying working memory and top-down attention processes, we trained macaques to match two patterns presented with a delay between them. As the above processes are likely to require communication between brain regions, and the parietal cortex is involved in spatial attention, we simultaneously recorded neuronal activities from the interconnected parietal and middle temporal areas. We found that mnemonic information about the first pattern was retained in coherent oscillating activity between the areas in high-frequency bands, followed by coherent activity in low-frequency bands that mediate top-down attention on the relevant location.

**ONE SENTENCE SUMMARY:** Gamma coherence allows retaining object features in a saliency map while lower frequency coherence facilitates attention.

---

Cognitive functions such as attention depend heavily upon communication between brain regions. Synchronised oscillations of memebrane potentials of groups of neurons in different areas have been postulated as a basic mechanism for the interaction between brain regions, a concept often referred to as ‘communication through coherence’, or CTC (*1, 2*, and illustrated in Figure 1A). If groups of neurons from two different areas have oscillations of similar frequencies it can help in transmitting signals from one area to the other by improving the chances to reach the threshold for initiating action potentials. The CTC has been demonstrated in macaques between parietal and an early visual area, namely the middle temporal area (V5/MT) (3), prefrontal and parietal cortices (4) and between prefrontal cortex and visual area V4 (5) in tasks requiring top-down attention. It has also been suggested (6–8) that attention-demanding tasks such as finding an object in a cluttered scene or matching a stimulus to an item in working memory may involve a two-stage process. At the first stage the most relevant spatial locations are selected based on coarse featural information from the visual scene, and at the second stage a spotlight of attention focuses attention on one of the selected location at a time for detailed visual processing and to enable the object of interest to be identified.

**Figure 1.**
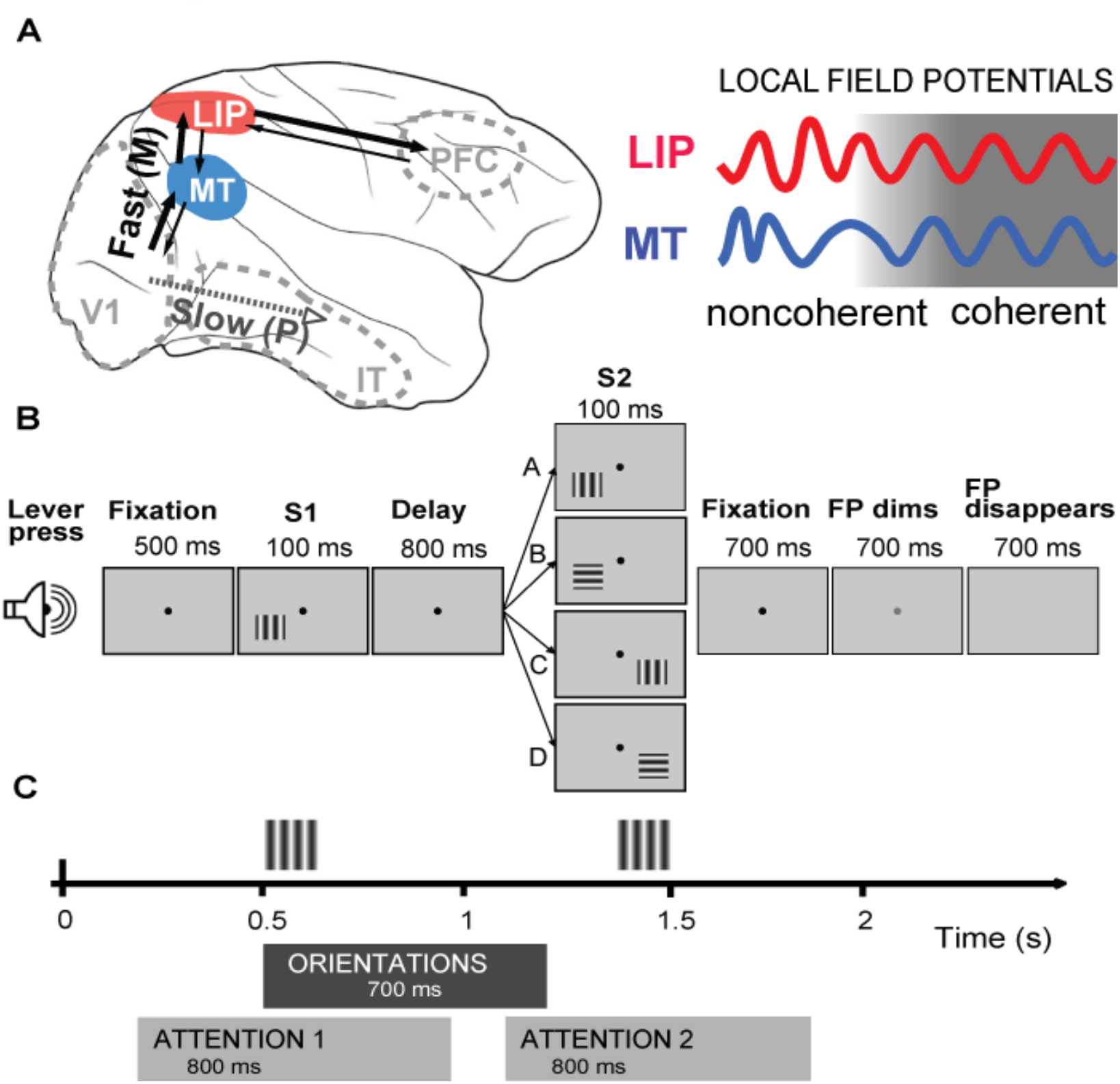
Recording areas in the monkey brain and the experiment paradigm. Areas MT and LIP within fast dorsal visual pathway are shown on the left, and two LFP signals on the right, showing initial lack (noncoherent) and subsequent presence (coherent) of synchronization. Fast dorsal pathway is believed to provide spatial information first in order to facilitate detailed recognition of the objects by the slower ventral pathway (7,8,9). (B) Schematic depiction of the DMS task. The task involved matching two gratings (S1 and S2) presented briefly for 100 ms each with an interval of 800 ms between them (Delay). The monkey presses a lever when ready for a trial and fixates on the central point (FP) during the trial. Gaze fixation was controlled by the infrared oculometer and the trial was aborted if gaze deviated by more than 1°. The monkey was required to report whether each trial was a match (both gratings appearing at the same location and having the same orientation, case A) or a non-match (whenever the locations or the orientations or both were different - cases B, C, and D). For match trials monkey had to release the lever when the FP dimmed, and for the non-match trails when it disappeared. (C) Time course of the match trial with the intervals used for coherence comparisons. Feature-related coherence bands were analysed by comparing coherences evoked by stimuli of 2 orthogonal orientations in 700ms long period after S1 onset (ORIENTATIONS), whereas attention-related coherence band were obtained by comparing 800ms intervals around S1 and S2 (ATTENTION1 and ATTENTION2).

Recent findings obtained in monkeys performing a Delayed Match to Sample Task (DMS, see Figure 1B), namely matching visual stimuli presented with a delay between them suggest that the lateral intraparietal area (LIP) of the macaque’s posterior parietal cortex may possess the neural substrate for such a two-stage process. This is particularly plausible given the presence of feature selective neurons, in LIP, e.g. cells with selective responses to differently orientated lines (9–12). We have also recently shown that LIP contains two groups of neurons having distinctly different responses to the stimuli in our DMS task, which occur during different parts of the delay period between the stimulus pair (12). One group of cells are feature selective and show higher activity in an early part of the delay period between the two stimuli in each trial, but not any attention-related elevation around the second stimulus of the pair (Attentional Enhancement negative, or AE− cells). The second group of cells (AE+) show poor feature selectivity, but exhibit enhanced responses around the second stimulus when attention is attracted to that stimulus location by the first stimulus. Area MT provides the major afferent input to LIP necessary for the feature selectivity expressed by the LIP. In turn, area MT receives top-down attention signals from LIP which increases responses of MT cells (3). However, since spiking activity of feature-selective cells in MT usually declines within 200 ms from stimulus offset (3, 13), transfer of the featural information by neuronal spiking ostensibly occurs within a short interval. Therefore in DMS taks, featural information that would drive the LIP AE+ cells for eliciting top-down attention later during the delay period is stored in some way other than as a simple increase in spike rate in both areas after the stimulus-evoked afferent signals from MT ceases. This question of the nature and site of the transition from coding feedforward feature information to coding feedback attention signals remains open. We suggest that synchronization of the oscillatory activities between MT and LIP may serve as the transition mechanism. CTC which mediates interareal communications may use different oscillating frequency bands in the bottom-up and top-down pathways (14, 15), and thus the transition might be accompanied by a change in the coherence frequency.

To address the above questions, we analysed local field potentials (LFPs) and spike trains which were simultaneously recorded from retinotopically matching sites of areas LIP and MT (n=36) while the monkeys were performing the attention demanding DMS task (Fig. 1B). As the monkey had to match stimuli by both location and the pattern, the monkey’s attention had to be directed to both the stimulus feature and where it was presented (Fig. 1B; see also supplementary materials). We had reported earlier that in this task, feedback LIP signals led to increased responses in topographically corresponding, attended locations in MT and reduced responses in unattended locations (3). Such spatial attention feedback was facilitated by coherent oscillations between the two areas in the beta-to- low gamma frequencies (25-45 Hz) during the late delay period and during the response to the second stimulus. The prominent characteristics of this LIP to MT feedback were its focal attention on the location without any feature selectivity and its reliance on the AE+ cells of LIP (3, 12), suggesting that the transfer of featural information from AE− to AE+ cells occurs earlier, requiring a separate process. Here, we first determined the range of LFP frequencies which showed significant feature-related or attention-related coherence between the two areas by applying the Aversen statistical technique to multiple comparisons across frequencies (16) (see Fig. 2A and Supplementary Materials for details). Fig. 2A shows frequency ranges sensitive to feature discrimination or to focal attention – namely the frequencies where significant coherence differences were found between conditions: feature-related for comparisons between features of stimuli or attention-related for comparisons between periods of low and high attentional load. Feature-related coherence differences were found in 17 pairs of recordings and attention-related coherence in 19 pairs. It can be seen that feature-related coherence tends to be in the higher gamma frequency bands (40-200 Hz, blue lines), whereas enhanced attention-related coherence is largely seen in the beta and low gamma frequencies (13-45 Hz, red lines), accompanied by suppression of coherence in high frequency range (above 47 Hz, grey lines).

**Figure 2.**
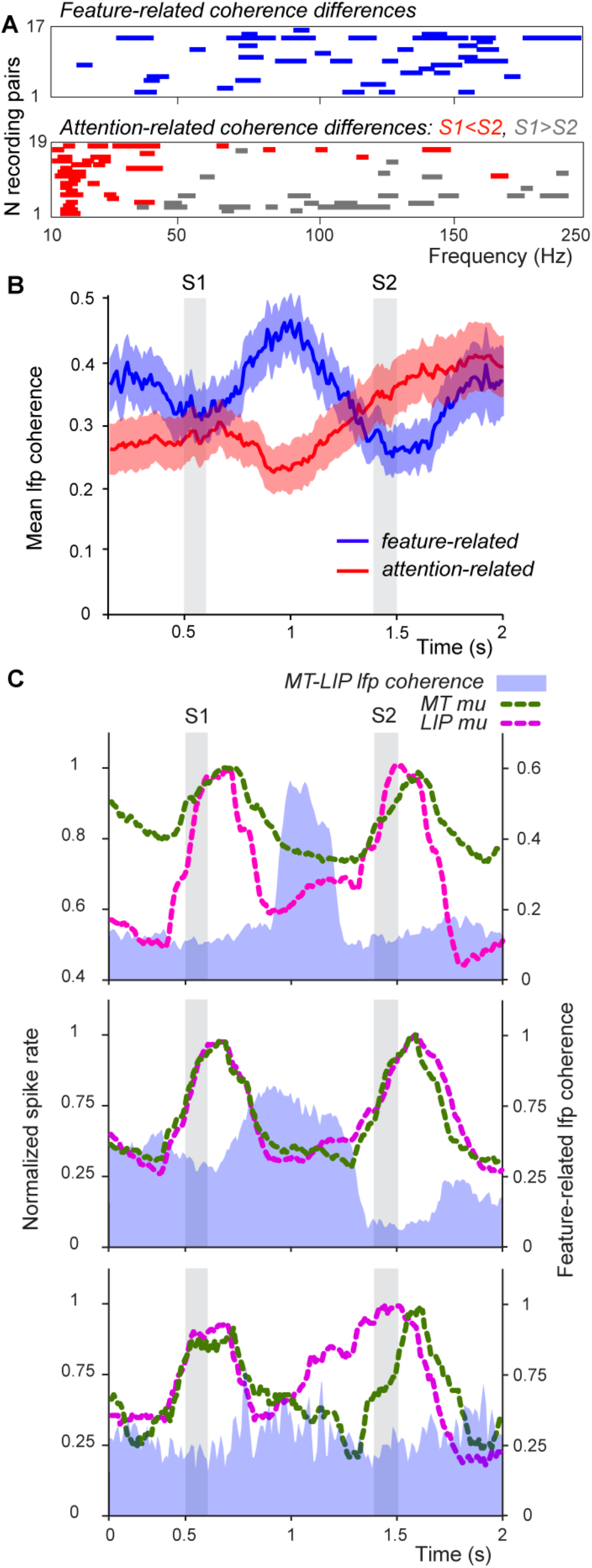
(A) Frequency bands where significant LFP coherence differences were observed. Feature-related differences (calculated using responses to the optimum and orthogonal orientations of the first (S1) grating, when attention has not been captured by the location or grating orientation) are shown in blue. Attention-related coherence differences (calculated from responses to grating of optimum orientation on RF as S2) are shown in red when coherence was enhanced in the Attention 2 interval (in match trials - case A and in grey when it was suppressed in the Attention 2 interval (in non-match trials - cases C and D). (B) Mean dynamics of significant LFP coherences averaged across all recording pairs with the same trial structure (N=14 for feature-related and N=17 for attention-related). Feature-related coherence dynamics is shown as dark blue line, attention-related (enhancement) as dark red line. Light-coloured areas represent ±SE. Grey rectangles (S1 and S2) designate periods of stimulus presentation. (C) Normalized multiunit response in LIP (purple dotted trace) and MT (green dotted trace) simultaneously recorded, showing mean spike rates for the match trials when both S1 and S2 had preferred the orientation. Each panel represents one pair of recording sites (MT-LIP). Spike rate is normalized to maximal response to S1, with scale on the left. Corresponding time course of feature-related LFP coherence is shown as the blue area, with its scale on the right.

We next analysed the temporal course of LFP coherence between MT and LIP sites just in the ranges of frequencies where either significant feature-related or significant attention-related coherence occurred. The blue trace in Fig. 2B shows the course of the feature-related LFP coherence between MT and LIP sites (n=17) which rises and reaches its maximum during the delay period, approximately 400 ms after the offset of the first stimulus, S1. Thereafter, it comes down to baseline level approximately 200 ms before the onset of the second stimulus, S2. On the other hand, attention-related coherence, as shown by the red trace (n=19), begins to rise only about 300 ms prior to S2, just as the feature-related coherence declines. For much of the course of the trial, there is also a general negative correlation between the two coherences (Pearson r= −0.54, p<0.001), when calculated over the whole length of the trial).

We also checked the possibility whether the feature-related coherence seen in high gamma frequencies could be caused by ‘spike leakage’, an artefact due to the LFP reflecting lower frequency components of action potential waveforms (18, 19). We did this by comparing the time course of the coherence in these significant gamma frequencies with the time course of multiunit responses (see supplementary materials for details). Fig. 2C shows the results for three of the recorded pairs. As can be seen in these three example cases and also when analysed across all those recording sessions where there was both significant LFP coherence and a multiunit response (MU) to the first grating, there was a significant latency difference between the spike response maxima and the coherence maxima in both areas (LIP: N=14 recording sites, p<0.001; MT: N=15, p< 0.001, Wilcoxon Signed Rank test). In MT, the peak spike response preceded the coherence maximum by 279 ms and in LIP by 341 ms. A recent study done in area MT also shows that synchrony seen at certain high gamma frequencies (180-220 Hz) represents sensory signals rather than a reflection of spike leakage (20), adding further credence to the importance of high gamma frequencies in information processing.

Our results suggest that the feature-related synchrony between MT and LIP in the early part of the delay and occurring after the vigorous response to the S1 stimulus is likely to be the first stage of a two-stage model of attention. Thus, the relevant feature information may be retained as a working memory buffer at a site where it can be readily used for directing top-down attention (21), consistent with the suggestion that area LIP constructs such a saliency or priority map for this purpose (22). To test this further, we analysed whether neuronal spiking was related to the amplitude of the LFP oscillations involved in featural coherence in the period of its maxima. This was done by calculating spike-triggered average (STA) of the amplitude envelopes of the oscillations occurring in the relevant frequency bands (Figure S2). The envelope helps to reveal the amplitude modulation of the oscillation and to relate neuronal spiking to the changes of that amplitude (Fig. S2A). Using multiunit spikes as the trigger we found that STAs in the majority of the recording sites in both MT (13/17) and LIP (17/17) were significant compared to pseudospike-triggered STAs (Details in Supplementary Materials, illustrated in Fig. S2B). Moreover, all AE+ cells derived from the same LIP sites as the envelope (n=9) triggered significant STAs, while only a small fraction of AE− cells had such a relationship (n=3/13), making AE+ cells likely recipients of the feature-related information retained during early delay period (p=0.0001, Exact Fisher test). This result is consistent with the suggestion that cortical synchronization may play a role also in working memory, especially in retention of sensory information for short periods (23, 24). Given the temporal sequence of the coherencies in our results, we propose that the feature-related LFP coherence drives the oscillatory activity among LIP’s attention-related sites, which in turn drives the oscillations in topographically corresponding sites in MT, as seen here and in our earlier study (3).

The presence of feature-related synchronised oscillatory activity in the feedforward MT to LIP connection during the delay may be the site of the earliest emergence of working memory in the brain, prior to its presence in neural activity within LIP itself (12) and its well-documented presence in the prefrontal cortex (11, 13). This is consistent with the findings suggestive of integration at the level of LIP of siganls from early visual areas, in particular from area MT (11). It is known that spiking activity in MT does not reflect working memory trace during the delay period (3, 13), while higher-order areas such as lateral prefrontal cortex (LPFC) possess neurons that are active during delay period (13). However since feature sensitivity appears in MT prior to these higher-order areas, our results suggest that the short-term retention of the sensory input prior to its being used for attention-related activity in the frontoparietal network may be in the form of coherent LFP oscillations between MT and LIP. Such coherence-dependent representation without elevation of neuronal firing rates in MT is consistent with the elevation of the LFP power observed in MT during the delay period in the absence of increase in the firing rates of individual neurons (13). Thus transient storage of featural information can be LFP coherence-dependent, while more sustained representation of the item in memory would be associated with spiking activity in higher-order visual areas. It is however an open question whether in our task, the transition from feature selectivity in the early delay period to attention related activity in the late delay period involves the prefrontal spiking activity or takes a more direct route within LIP, but with some degree of prefrontal modulation.

The two sets of coherent oscillations we found that operate at two different ranges of frequencies, one at beta/low gamma and the other at higher gamma, demands both a mechanistic explanation as well as a possible functional role. The biophysical characteristics of the cells and the circuitry they are embedded in leave them with a range of frequencies that display optimal resonance (26). Thus it is plausible that the feature-selective oscillations happening in LIP’s input layers arise from cells with a particular morphology, possibly smaller stellate cells, whereas cells in the deeper, output layers providing feedback to MT are larger pyramidal cells, which, due to their lower input resistance and longer time constants are likely to have a lower resonance frequency. Since the purpose of an attentional feedback to early visual areas is likely to be to select the object location for more detailed processing of all features associated with the object (7, 8), it is more meaningful for the feedback to be simply spatial and acting on all feature domains at the object locus. In fact, the LIP feedback to MT seems to be purely spatial and makes little distinction between the different feature-selective cells in the corresponding retinotopic location (3). Human psychophysical experiments also reinforce the point that gating functions of the dorsal stream prioritise spatial locations rather than specific features (27). The different frequencies for the two functions may also be functionally fortuitous, consistent with recent evidence that the brain may use different synchronising frequencies for different functions (20).

In conclusion, our findings suggest that feature-selective coherence may represent object-related information stored in a saliency map which is used to drive the activity of the cells directing spatial attention. Such a system would enable not only serial allocation of attention as in classical visual search situations (12), but also for deliberately delayed responses after a stimulus presentation, since external featural information may not be continuously available due to occlusion of objects or due to saccades made to different locations in the visual scene.

## Acknowledgements

We thank Dr. Ivan Pigarev for taking part in some of the early studies. We are grateful to Drs. Andrew Metha and Chris French for critical comments on the manuscript.

## Author contributions

Y.S. and T.R.V. conceptualised and performed the original experiments and collected the data. E.L., T.R.V. and M.K. conceptualised the present model. E.L. developed the analytical tools for studying the model. E.L. and M.K. did the bulk of the new analysis. E.L. and T.R.V. wrote the original draft. All authors critically reviewed and edited the final manuscript. T.R.V. acquired funding and administered the project.

## Data and materials availability

All data that are not presented in main text or Supplementary Materials, are available from the corresponding author upon reasonable request.

## Supplementary Materials

### Materials and Methods

#### Animal care and behavioural training

Data was collected from two male macaque monkeys (*Macaca nemestrina)*. The study was conducted as per the guidelines of the National Health and Medical Research Council Australian Code of Practice for the Care and Use of Animals for Scientific Purposes and approved by the University of Melbourne Animal Experimentation Ethics Committee. Monkeys were housed together and had ad lib access to water. They were trained to come voluntarily to the training chair and perform a visual delayed match-to-sample task (DMS, Fig 2B). For detailed description of husbandry, surgical and training procedures, please refer to Supplementary Online Material of (3).

The task included presentation of two sinusoidal gratings (8^0^×8^0^, 30% contrast with a mean luminance of 15 cd/m^2^) with an interstimulus interval (ISI) between them. The monkeys had to match two gratings with respect to both the orientation and location of the gratings. The orientation preference of neurons at the recording site was first assessed by hand held stimuli, then verified with gratings, before each set of recordings; and an orientation close to that preferred for multiunit activity in both MT and LIP, and the orientation orthogonal to the preferred, were used for the recordings. Up to 5 different locations in the visual hemifield around the fixation point and the two orientations were presented in pseudorandom succession in such a way that approximately 50% of the trials were ‘Match’ trials, where the second grating appeared at the same location as the first and the orientations of both gratings were also the same. The pseudorandomisation was also mildly biased for the first stimulus (S1) to fall on the receptive field location of the MT and LIP recording sites to ensure collection of a sufficient amount of data from the superimposed receptive field sites in the two cortical areas.

The monkey initiated each trial by pressing a lever. This led to the black fixation spot (FP, 0.1° diameter) being presented at the centre of a uniform grey screen (of luminance 15 cd/m^2)^. The monkey was required to maintain fixation during the whole trial, but could break it between trials. If an eye movement of more than 1°, monitored using an infrared oculometer (Dr. Bouis) was detected, the trial was aborted.

S1 was presented for 100 ms duration, 500 ms after the start of fixation. After a delay period, the second grating (S2) was also presented for 100 ms duration. The delay was set for 800 ms in 29 recordings, 900 ms in 1 recording, 1000 ms in 1 recording and 500 ms in 5 recordings. The monkey had to keep fixating for an additional 700 ms after S2 offset. After that, the FP was dimmed for 700 ms before it disappeared. In Match trials (when locations and stimulus orientations of S1 and S2 were the same) the monkey was required to release the lever within the 700 ms dimming interval to obtain fruit juice reward. In all Non-Match trials, the lever had to be released only after disappearance of the FP. Correct responses were rewarded with fruit juice and incorrect responses were followed by prolongation of the ISI by 1-3 seconds. The window allowed for response was from 200 to 650 ms after either the start of the dimming period (in the case of match trials) or after FP disappeared (in the case of non-match trials).

#### Electrophysiology

A low invasive “halo” surgical technique was used for implantation (3, 28, 29). Structural MRI was performed prior to the start of experiments to guide craniotomies (2.5 mm diameter) necessary to record from LIP and MT areas.

We recorded neuronal activities from areas LIP and MT using tungsten or platinum– iridium microelectrodes (FHC, ME USA). The recorded signal was filtered either at 1–4000 or 10–4000 Hz and sampled at 10,000 Hz by the Cambridge Electronic Design (CED) Micro1402 data collection system. It was further filtered using inbuilt filters of CED’s Spike 2 programme to obtain separate spike and local field potential (LFP) data. A band-pass filter was applied in the 300 – 4000 Hz range for spikes and a low-pass filter up to 250 Hz for LFPs. Single or multiunit activity was acquired with the WaveClus spike sorting toolbox running on Matlab (30) with amplitude threshold set to be above 3 standard deviations from baseline noise of the band-pass filtered data.

36 paired LIP-MT recordings with overlapping receptive fields and matched preferred stimulus orientations were used for the present analysis.

#### Data analysis and results

##### General approach to coherence analysis

Coherence analysis allows one to measure synchronisation between two signals, which in our case, was the degree of synchronisation between the neuronal activities recorded simultaneously from areas MT and LIP. Since neuronal activity can potentially occupy a large range of frequencies, it is necessary to define an approach to select specific frequency bands for further analysis.

The usual way to analyse neuronal activity in the frequency domain largely depends on assumptions regarding putative functions of the oscillations in different frequency bands. To overcome such semi-arbitrary character of choosing bands of interest and the questionable applicability of widely defined bands to activities of different cells with different soma sizes, axonal arbours and synaptic characteristics, we compared coherences by applying Aversen’s technique to multiple comparisons across frequencies, implemented as “two_group_test_coherence” function in Matlab-based Chronux toolbox (http://chronux.org/, 31, for the details see 16). This approach permits splitting the analysis into two parts where the first one addresses the relevance of a particular oscillation to the behavioural task and the second step involves analysing the data selectively just in those bands that were found relevant to observe dynamics of coherence change along the trial specifically within the task-relevant frequency bands. At both steps, we applied multitaper spectral estimations implemented in Chronux toolbox.

Detailed description of the coherence comparison can be found in (16). Here we describe only the details relevant to the application of this method to our data.

The coherence is calculated as:

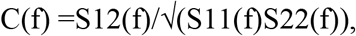

where S(f) is the spectrum, 1 and 2 refer to the neural activity in MT and LIP respectively. Coherence was calculated involving 3 orthogonal Slepian taper functions and a time bandwidth product of 2 (16, 32, 33). For data length N and frequency bandwidth W, the first *K* = 2*NW*−1 Slepian sequences are optimally concentrated in the frequency range [−*W*, *W*]. Therefore the minimal frequency range for estimating significance of the coherence differences between 2 conditions is

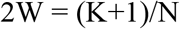

The method of comparison of coherences is based on the fact that the Fisher *transformed coherences*, tanh^−1^(*C*(*f*)), has Gaussian distribution and therefore differences of transformed coherences for two populations are distributed as Δ*z*(*f*)~*N*(0, 1) when the two population coherences are equal.

Statistical estimation of the coherence difference utilizes jackknifed variance based on computing *m* = *N(trials)K(tapers)* Fourier transform pairs, leaving out one taper and one trial in turn. To summarize, coherence differences (Δ*z*(*f*)) are considered significant if they exceed jackknife-based error bars consecutively for at least 2W frequencies. We report here only those differences that exceeded these criteria.

Figure S1 illustrates oscillatory behaviour of LFPs in areas MT and LIP in one example Match trial taken from a recording session where multiunit (MU) activity in response to S1 and S2 were significantly above the baseline and significant coherence between the two sites was observed. Note that high gamma oscillation was evoked by the first stimulus and lasted over approximately half of the delay period in both MT and LIP and thereafter replaced by lower frequency oscillations. We applied coherence analysis to this specific pattern of activities.

##### Feature-related coherence differences

The two conditions that were compared to estimate feature-related activity were presentations of two orthogonal orientations of the grating, each presented as the first of the pair in all trials, where S1 fell on the receptive fields of the neurons in the two sites.

That first grating could appear in any of 5 different locations meant that attention was not already focussed at any one location at the time of S1. Furthermore, the monkey had to code in memory the location and the orientation of S1 grating to be able to match them with those of S2. Therefore, since the location corresponding to the RF does not change, any difference in neural activity between the two orientations in the period after S1 onset, including that during the immediate delay period following S1 can be assumed to be orientation-related activity. Our previous results also indicate that feature-related neuronal activity in LIP can indeed be retained for some time during delay period but ceases towards the end of it (3, 12). Therefore for the recording sessions where delays were equal or more than 800 ms (N=31), we decided to choose a 700ms interval from S1 onset for the first step of our analysis, as it included S1 presentation and a reasonably long interval after that (600ms), but not the later part of the delay period (last 200ms). Coherence for all trials where one orientation was presented as S1 were compared to the coherence for all trials with the orthogonal orientation of S1 using the above described statistical technique (16). Thus for the 700 ms interval used for feature-related activity estimation, the 2W frequency bin was equal to 5.71Hz. Thus coherence differences exceeding jackknife-based error bars for more than 5.71Hz were taken as feature-relevant. The 2W value was adjusted according to the same principle for the recordings with 500 ms delay period (N=5) where the interval of the analysis was reduced to 500ms starting from S1 onset. The second step of the analysis involved constructing the dynamics of the feature-related coherence. Coherence was calculated in 300 ms intervals in 10 ms steps along the trial. Since direct estimation of orientation preference in the frequency domain is problematic, we used as preferred the orientation that produced more widespread changes in terms of frequency bandwidths that were deemed relevant in step 1. To visualize the dynamics of coherence, coherences for all relevant frequency bands were averaged within each recording site for that orientation. For visualization purposes only Match trials are used because responses to different S2 gratings as in non-match trials can be dramatically different (3, 12)

Latency of the coherence maximum within the interval of interest (700 ms or 500 ms after S1 onset) was later compared to the corresponding latencies of the multiunit responses in MT and LIP in the same interval using *Wilcoxon Rank-Sum test.*

##### Attention-related coherence differences

In the DMS task, covert attention is required to match location and orientation of the second stimulus to the first one. Attention to the S1 location in order to match S1 to S2 creates ramping of the spike rates in the late delay period and enhanced response to S2 (3). To estimate attentional modulation we compared activity around S1 to that around S2 for trials with matching S1 and S2. As described before, attentional modulation starts before S2 onset in the late part of the delay period (3, 12) and finishes after S2 offset. Therefore, to cover the whole interval we chose two 800ms windows, one around S1 and another around S2, from 300ms before stimulus onset to 400ms after stimulus offset. For the chosen 0.8 sec interval used for attention-related activity estimation, the minimal 2W frequency range for significant coherence differences is 5Hz. As short delay periods are known to show an attentional blink in LIP responses (34), we included in this analysis only those recordings where delay period was set to 800, 900 or 1000 ms (31 recordings).

Increased coherence during S2 was considered as a sign of the positive attentional modulation occurring in a particular frequency band, while smaller coherence values during S2 were regarded as negative attentional modulation. Positive modulation indicates frequency bands where coherence is enhanced in order to sustain attention, while negative marks the processes that are suppressed. For the second part of the analysis, which aimed to show the temporal dynamics in the frequency bands identified as attention-related, we used the same window parameters as for the dynamics of feature-related coherence differences, namely 300 ms intervals in 10 ms steps along the trial. We analysed positive S2>S1 (attentional enhancement) and negative S2<S1 (suppression) coherences separately.

To visualize coherence dynamics, coherences for all significant frequency bands were averaged within each recording site.

##### MU responses in MT and LIP areas in relation to feature-related coherence

Since feature-related activity occurs in the high frequency range where the LFP can be potentially affected by leakage of spiking-related frequencies (18, 19), it was necessary to test if the observed LFP coherence can be simply explained by the higher spike rate from the response to the visual stimulus. Spike removal from the LFP signal may not be particularly effective in the case of extracellular recordings made with relatively low impedance electrodes. Therefore we compared latencies of the response peak of MU to the latencies of the peak LFP coherence *as the spike leakage can be expected to be maximal for the maximal spike rate.* For every recording site, we identified the presence of the response to S1 by comparing mean background spike rate in 300 ms before the S1 onset to mean spike rate within 300 ms after S1 onset using Wilcoxon Signed Rank Test. Recording sites with significant S1 response to any of the two orientations (*p* < 0.05) were used for further analysis. In order to investigate if the coherence maximum corresponds to the MU maximum, latency of the S1 response maximum was estimated using the same averaging window and step parameters as those used for analysis of coherence dynamics, namely 300 ms window duration in 10 ms steps. Latencies of MT and LIP MU response maxima were compared to the latency of the LFP coherence maxima using the *Wilcoxon Rank-Sum test, for the trials with significant feature-related coherence.*

Another possibility to consider is that the activity of other cells that do not respond to the stimulus may be active in the delay period and contribute to the coherence seen in the delay period. This synchronized activity is best observed in the multiunit response rather than in single unit recordings, especially in the case of higher oscillation frequencies, because an individual cell that is in phase with an oscillation may not fire action potentials with every cycle (35). Therefore we also studied spike-spike coherence dynamics between multiunit spiking activities in MT and LIP areas for feature-related processes in order to compare it to the LFP-LFP coherence, applying again the same parameters as for the LFP analysis described earlier. The spike-spike coherence difference between the two orientations reached significance only in two of the recording pairs. However it is to be admitted that that spike-spike coherence is generally weak as action potentials do not accompany each cycle of LFP oscillation, resulting in difficulties in identifying spike-spike coherence, especially in the high frequency range.

##### Spike-triggered average (STA) of feature-related oscillations with multiunit activity as the trigger

It has been suggested that interareal communication may not directly evoke responses of cortical neurones, but may just change their excitability by inducing coherence between the two sites (35). This has indeed been observed in the attentional modulation of MT by LIP in the delay period just prior to the attentional enhancement seen in the visual response (3). A saliency map relying on initial featural input according to this view can be seen as a “patchwork” of synchronized and desynchronized cell assemblies. As synchronised input is more likely to elicit spikes, we analysed the amplitude modulation of the coherent oscillations and the dependency of spiking to that amplitude modulation.

The LFP was band-pass filtered in those frequency bands where we observed significant feature-related coherence difference, using bidirectional Butterworth filter to achieve zero phase-shift. The filter had just 3 poles to avoid ripples when band-pass filtering using relatively narrow frequency bands. Instantaneous amplitude of the oscillation – the envelope of a filtered signal – was obtained by calculating the Hilbert transform of the detrended and filtered signal and using its complex modulus (magnitude) as shown in Figure S2A. The resulting signal provides information regarding the amplitudes of the oscillation in question, and the possibility to test whether the cell spikes are related to the amplitude modulation of that oscillation. In each of the relevant frequency bands, STA was calculated for both the oscillation and its envelope using multiunit spikes as triggers and z-scored oscillation/envelope signals. z-scoring is done by subtracting the mean and dividing the signal by its standard deviation in order to normalize all signals to one scale. For every recording site, we compared the peak-to-trough amplitude of the averaged signal wave triggered by multiunit spikes to the distribution of the analogous values obtained by taking randomly the same number of pseudospikes 1000 times. The peak-to-trough amplitude was calculated within the interval covering one cycle of the averaged oscillation. STA was considered significant if its peak-to-trough amplitude was above 95% of the randomly taken STAs triggered by pseudospikes. The spikes used to compute STAs occurred in the early delay period (from 300 to 700 ms after S1 onset) when the feature-related coherence maximum was observed and when the multiunit response to S1 had already faded.

##### Spike-triggered average of feature-related oscillations (STA) with individual AE+ or AE− cells activity as the trigger

The STAs were calculated in the same way as described in the previous section, but using as triggers the spiking activities of each of the AE+ and AE− LIP cells recorded from the sites which exhibited significant feature-related LFP coherence difference (13 AE− and 17 AE+ cells). The AE+/AE− classification of a particular LIP cell was based on the Attentional Enhancement Index,

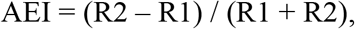

where R1 and R2 are responses to stimuli S1 and S2 in match trials and both stimuli are presented within the receptive field of the cell (for the details see 12).

---

**Figure S1.**
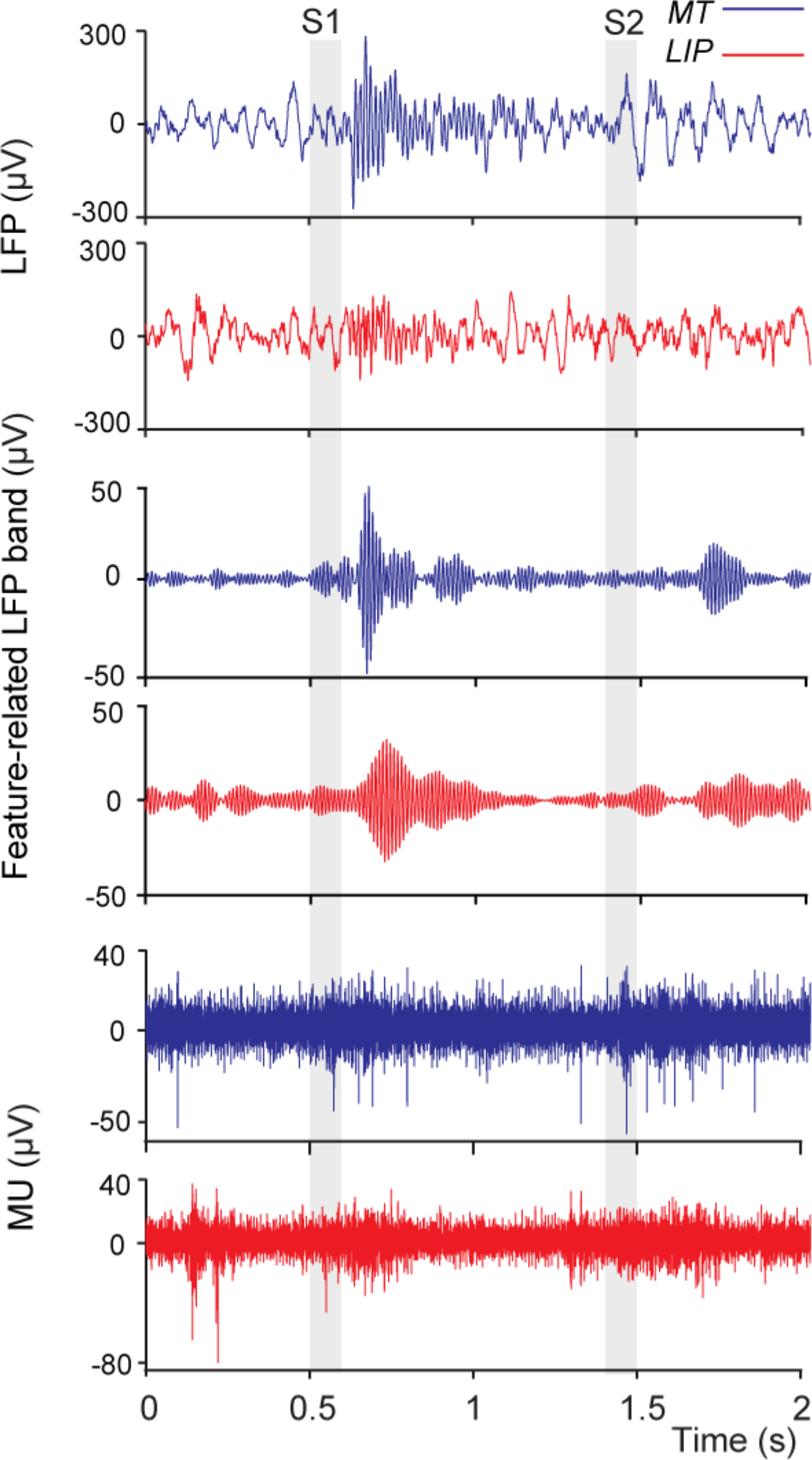
LFPs recorded from MT and LIP areas during a single match trial (upper panel), band-pass filtered LFP where all but the coherent gamma frequencies are filtered out (middle panel), and the corresponding MU responses (bottom panel). Gray intervals correspond to S1 and S2 presentation.

**Figure S2.**
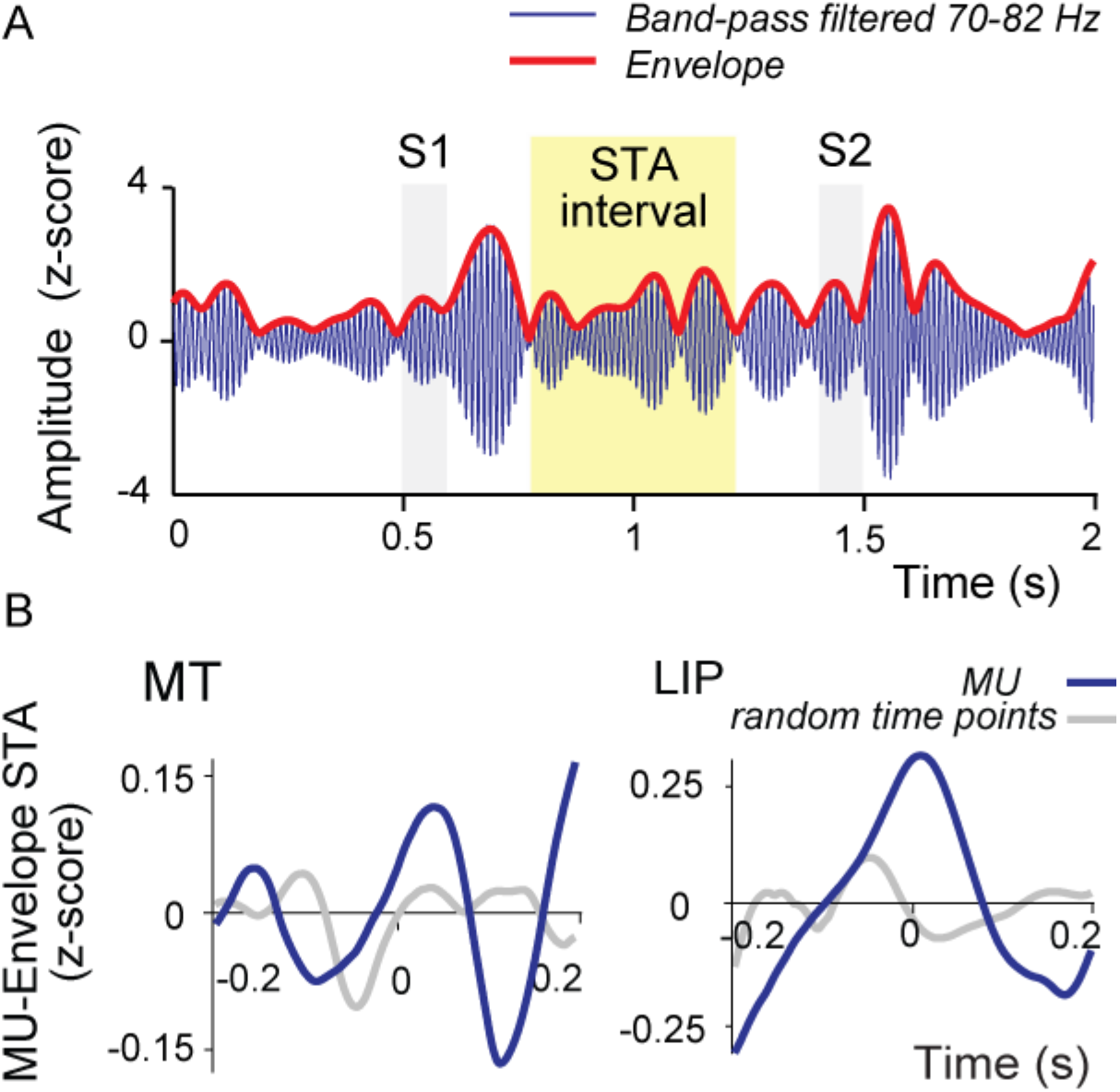
(A) Example of LFP during a single trial, transformed for STA analysis. Blue trace represents LFP filtered in the frequency range where significant feature-related coherence difference was found and the red trace shows the magnitude of Hilbert transform of the filtered LFP (Envelope). Gray intervals correspond to S1 and S2 presentations and yellow area shows the interval used to calculate STA. (B) Examples of STAs calculated with MU simultaneously recorded in MT and LIP as triggers and oscillation envelope as the averaged signal. STAs are shown for the recordings made with S1 being of the preferred orientation. Dark blue lines represent STA triggered by MU, gray lines STA triggered by pseudospikes.

